# Robot-aided Training of Propulsion During Walking: Effects of Torque Pulses Applied to the Hip and Knee Joints During Stance

**DOI:** 10.1101/2020.05.26.117036

**Authors:** Robert McGrath, Barry Bodt, Fabrizio Sergi

## Abstract

The goal of this study is to evaluate the effects of the application of torque pulses to the hip and knee joint via a robotic exoskeleton in the context of training propulsion during walking. Based on our previous biomechanical study, we formulated a set of conditions of torque pulses applied to the hip and knee joint associated with changes in push-off posture, a component of propulsion. In this work, we quantified the effects of hip/knee torque pulses on metrics of propulsion, specifically hip extension (HE) and normalized propulsive impulse (NPI), in two experiments. In the first experiment, we exposed 16 participants to sixteen conditions of torque pulses during single strides to observe the immediate effects of pulse application. In the second experiment, we exposed 16 participants to a subset of those conditions to observe short-term adaptation effects.

During pulse application, NPI aligned with the expected modulation of push-off posture, while HE was modulated in the opposite direction. The timing of the applied pulses, early or late stance, was crucial, as the effects were often in the opposite direction when changing timing condition. Extension torque applied at late stance increased HE in both experiments range of change in HE: (1.6 ± 0.3 deg, 7.7 ± 0.9 deg), *p* < 0.001). The same conditions resulted in a negative change in NPI only in the single pulse experiment — change in NPI for knee torque: −2.9 ± 0.3 ms, *p* < 0.001, no significant change for hip torque. Also, knee extension and flexion torque during early and late stance, respectively, increased NPI during single pulse application — range of change in NPI: (3.4, 4.2) ± 0.3 ms, *p* < 0.001). During repeated pulse application, NPI increased for late stance flexion torque — range of change in NPI: (4.5, 4.8) ± 2 ms, *p* < 0.001), but not late stance extension torque. Upon pulse torque removal, we observed positive after-effects in HE in all conditions. While there were no after-effects in NPI significant at the group level, a responder analysis indicated that the majority of the group increased both NPI and HE after pulse application.

## I. Introduction

In recent years, robot-assisted gait training (RAGT) has been devised and implemented in both research and clinical settings. Currently, the majority of RAGT devices, designed specifically to rehabilitate gait, utilize one of the various controller forms (e.g., force control, position control, or impedance/admittance control), and controller update methods (e.g., assist-as-needed control, inter-limb coordination, or finite state machine), to ultimately promote specific features of gait kinematics [1]. Despite the convergence of research on these methods, the efficacy of RAGT based on kinematic control has not yet exceeded that of conventional gait therapies [2], [3]. The limited efficacy of these methods could be due to their lack of targeting specific functional mechanisms of gait, which are only partially described by joint kinematics.

Gait speed (GS) is a primary indicator of walking function in rehabilitation, as it is associated with a better quality of life [4]. To increase GS during walking, it is necessary to increase the anterior component of the ground reaction force, and specifically its time integral, referred to as propulsive impulse [5]. Propulsive impulse is generated during late stance, right before push-off. Propulsive impulse is determined by push-off kinetics and kinematics; specifically by the moment applied by ankle plantarflexor muscles, and by the posture of the limb during push-off [6]. An established metric for quantifying push-off posture is the trailing limb angle (TLA), defined as the angle between the line formed by the foot center of pressure and hip joint center, at peak propulsive force, and the vertical laboratory axis [7]. In healthy control participants and post-stroke patients, it was observed that the increase in TLA contributes more than the increase in ankle moment to the resulting increase in propulsive force [7], [8]. As such, both push-off posture and propulsive impulse are possible targets for modulation of propulsion during robot-aided training. While it is known that push-off posture and propulsion are associated, it is unclear how to apply a robotic intervention to modulate propulsion, and whether a robotic intervention modulating propulsion during training will ultimately influence walking after training.

As such, we sought to develop a controller capable of training propulsion during walking. We began by investigating the changes in joint moments associated with the experimentally imposed factorial modulation of GS and TLA [9], [10]. We then approximated the effects of push-off posture on joint moments with brief pulses of joint torque. We compiled the amplitude and time of the resulting torque pulses to analyze patterns associated with TLA modulation for each joint. We observed that at the knee, an increase in TLA was associated with extension torque in early stance and flexion torque in late stance. At the hip, early and late stance extension torques were associated with an increase in TLA.

In this work, we seek to determine whether the modulation in joint moments associated with a change in propulsion dynamics, can be applied by a robotic exoskeleton in the form of torque pulses to the hip and knee joints to modify propulsion dynamics in healthy participants.

First, we wished to investigate the instantaneous effects of single-pulse torque intervention on the kinematics and kinetics of gait. As such, for the first protocol of this study, we formulated a set of sixteen conditions of torque pulses, based on the torque patterns associated with TLA modulation in our previous study [9], [10]. The protocol consisted of applying torque pulses to the hip and knee joints of healthy individuals during single strides, using a lower extremity exoskeleton, the ALEX II, while they walked on an instrumented treadmill. We observed the effects of torque pulses on the outcome measures of hip extension angle (HE) and normalized propulsive impulse (NPI) and determined the predominant factors influencing these measures at the stride of pulse application and the following strides. We hypothesized that each pulse torque condition would modulate HE and NPI in the same direction as the expected modulation of TLA, for the stride of application and the following three strides.

Second, we wished to measure how subjects would adapt to pulsed torque training, and how they would respond right after exposure to training. As such, for the second protocol of this study, we selected a subset of eight torque pulse conditions with the largest effects on HE and NPI observed in the first protocol. The protocol consisted of applying the selected eight pulse conditions to the hip and knee joints for consecutive sets of 200 strides with the ALEX II before and after subjects were walking with the exoskeleton controlled to display minimal interaction forces. We hypothesized that each torque pulse condition would exhibit short-term adaptation to pulse application and de-adaptation following pulse application removal. Furthermore, we hypothesized that the outcome measures would exhibit the same direction of modulation during intervention as the single-pulse experiment.

## II. Methods

### A. Study Participants

The single-pulse experiment included sixteen healthy adult participants (13 male, 3 female) of age (mean ± std) 25 ± 2 yrs, height 178 ± 5 cm and mass 75.6 ± 8.5 kg. The repeated-pulse experiment included sixteen healthy adult participants (7 male, 9 female) of age 24 ± 3 yrs, height 171 ± 10 cm and mass 70.3 ± 16.7 kg. Three participants (1 male, 2 females) were common to both experiment. Participants were only included if naive to the purpose of the experiment and free of neurological and orthopedic disorders that would affect normal walking function. All participants gave informed consent according to the IRB protocol number 929630 at the University of Delaware and wore their own comfortable lightweight athletic clothing.

### B. Equipment

Data collections were conducted on an instrumented split-belt treadmill (Bertec Corp., Columbus OH, USA) that measured analog force/torque data. The ALEX II robot [11], a powered unilateral lower extremity exoskeleton, as seen in Fig. 1, was utilized to apply torque pulses about the right knee and hip joints of participants. The exoskeleton is suspended by a mobile carriage over the instrumented treadmill and secured from moving during experimentation by locking casters. A custom real-time controller written in MATLAB & Simulink (MathWorks Inc., Natick MA, USA) acquired signals from the instrumented treadmill and ALEX II and sent command signals to the two motors at a frequency of 500 Hz. The ALEX II contains two Kollmorgen ACM22C rotary motors with integrated Smart Feedback Devices (Danaher Corporation, Washington D.C., USA). These provide an emulated encoder resolution of 4096 pulses per revolution providing an effective hip and knee angle resolution of 4.4 × 10^−4^ deg.

**Fig. 1:**
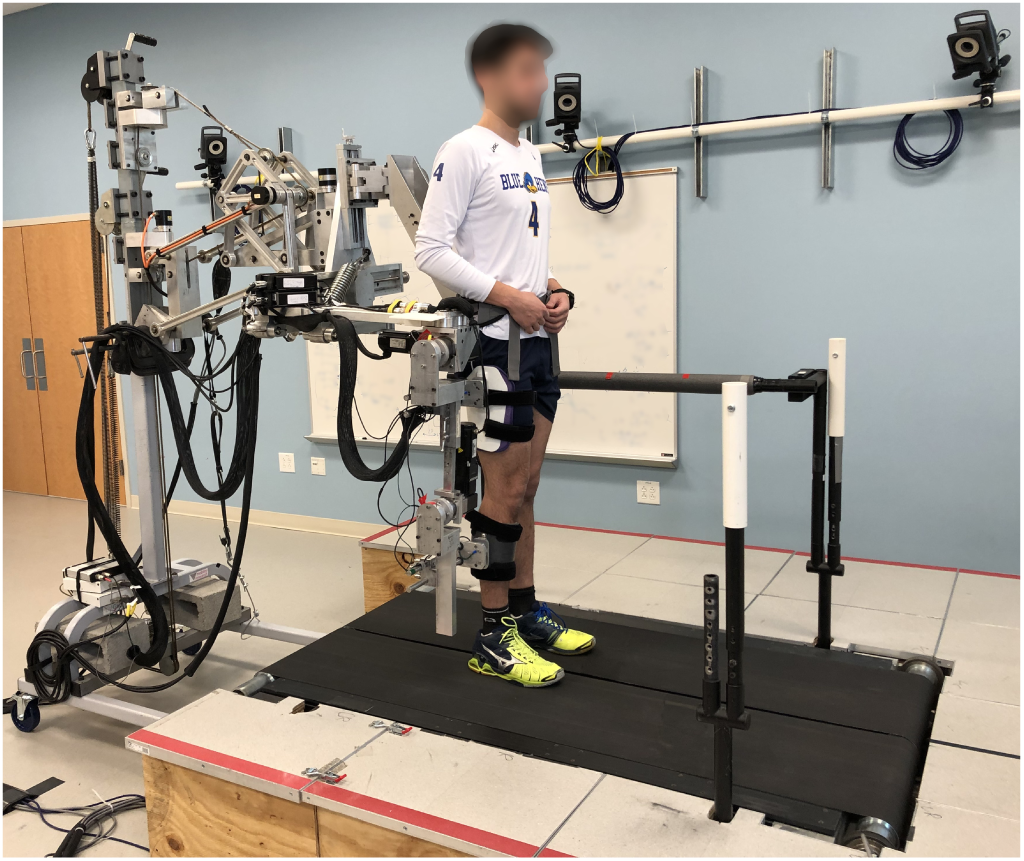
The experimental setup consisting of a subject in the Active Leg EXoskeleton II (ALEX II) while on the instrumented split-belt treadmill.

### C. Controller

A high level controller utilizes the vertical ground reaction force, as measured by the instrumented treadmill, to determine the time of right heel strike events and send desired torque values to the low-level controller. Heel strike events define the onset of the gait cycle and the time between events define the duration of the gait cycles. The average of the previous six gait cycles yields an estimated gait cycle time which is used with pulse time values, as percentages of the gait cycle, to determine controller event timing. The high-level controller’s selection of pulse condition, as per experimental protocol, specifies the joint torque amplitudes, onset as percentage of gait cycle, and duration as 10% of gait cycle.

A low-level torque controller featuring gravity and friction compensation was developed utilizing direct force feedback from two 6-axis force/torque sensors located between the exoskeleton structure and the shank/thigh cuffs. The implementation of force/torque sensor feedback allows for partial compensation of the high friction present in the geared motors and the inertia of the exoskeleton structure [12]. Section V-C of the Supplementary Materials describes the controller compensation for the delay in torque pulse application and a comparison between prescribed torque pulses and resulting joint torque.

### D. Experimental Procedures

#### 1) Single-pulse Experiment

In this experiment, torque pulses were applied only at one stride, followed by 5-7 strides of no pulse application to measure participants’ response. Sixteen pulsed torque conditions were tested in this experiment, as shown in Fig. 2. Each condition consisted of square waves with a duration of 10% gait cycle, and a timing of early or late stance, initiating at 10% or 45% gait cycle, respectively. The amplitude of these square wave pulses are 10 N·m or -10 N·m for knee extension or flexion, respectively, and 15 N·m or -15 N·m for hip extension or flexion, respectively.

**Fig. 2:**
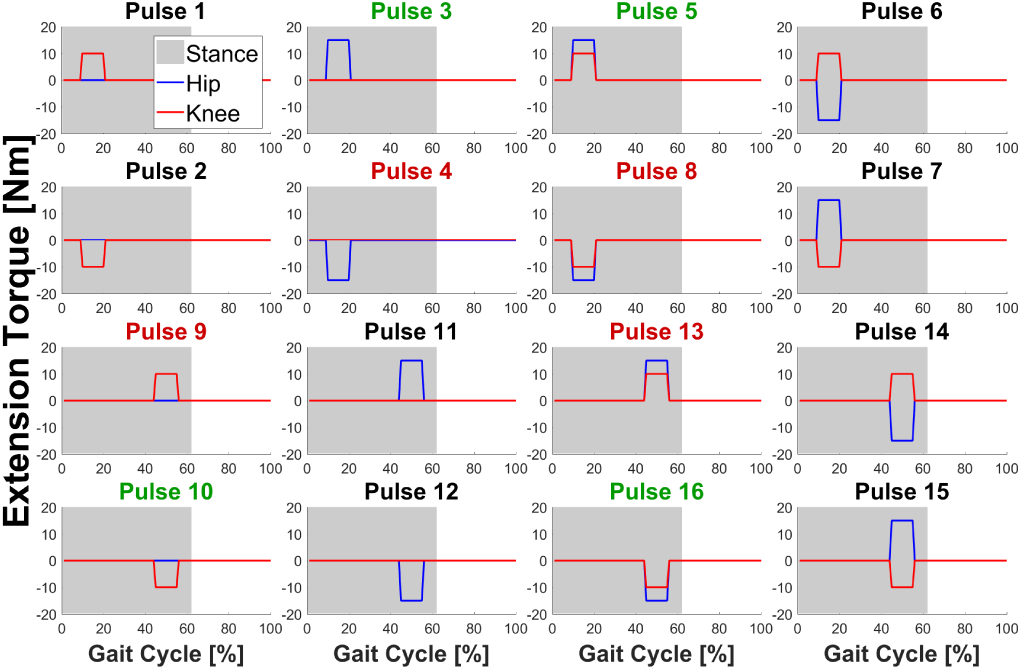
The 16 pulses were conditions implemented in the single-pulse experiment and the 8 pulse conditions, with colored titles, were implemented in the repeated-pulse experiment.

All participants performed two separate sessions, all while walking at their predetermined self-selected gait speed. Each session consisted of 5 repetitions of each of the 16 pulse conditions in a pseudo-randomized order. Both sessions featured the same sequence of pulses, lasted for a duration of 8-10 minutes, and were separated by a minimum of 5 minutes of rest outside of the exoskeleton. The participants walked at their self-selected speed, determined before the experiment while wearing the exoskeleton in zero-torque mode. Self-selected speed was determined and exoskeleton fitting was performed as in our previous work [13].

#### 2) Repeated-pulse Experiment

In this experiment, torque pulses were applied for 200 consecutive strides, preceded and followed by 100 strides of no pulse application, to measure participants’ response. A subset of 8 pulse conditions, out of the full set of 16 pulse conditions, were selected for testing in the repeated-pulse experiment (Fig. 2). The four pulse conditions with green and red titles increased and decreased, respectively, NPI in the single-pulse experiment. The selection process of these pulses is explained further in Sections II-E2c & III-A3

Each session consisted of a total of 700 strides in which strides 101-300 and 401-600 each consisted of a different pulse applied in consecutive strides. The first, middle, and last 100 strides were performed in the absence of pulse application, to allow for measurement of baseline and effects for each pulse condition. All participants performed a total of four separate sessions, two on each of the two days of experimentation while walking at their self-selected gait speed. Each session was separated by a minimum of 5 minutes of rest outside of the exoskeleton and lasted for 12-14 minutes. Each day of experimentation contained a grouping of 4 pulses with the same directional effect on NPI. Both the sequence of day (exposure to pulses increasing or decreasing NPI), and the sequence of the pulses within each day were pseudorandomized across participants.

### E. Data Analysis

#### 1) Outcome Measures

Two outcome measures were selected to describe the effects of the intervention on walking kinematics and kinetics. For kinematics, we selected hip extension angle (HE), as the angle of the hip of the right (robot-assisted) leg at the instant of peak anterior ground reaction force (GRF). For kinetics, we selected normalized propulsive impulse (NPI), defined as the integral of the anterior-posterior component of GRF over the time interval that the component is positive. Propulsive impulse was then normalized by the participant’s body weight (in N). Changes in outcome measures reported as percentages are normalized by the average baseline value of that outcome measure of the respective experiment.

#### 2) Single-pulse Experiment

In the single-pulse experimentation, outcome measures were observed at a total of five strides per applied individual pulse. This includes the prior baseline stride (−1), the stride of (0), and three strides following (1, 2, 3) application of torque pulses.

##### a) Linear Mixed Effect Models

We used linear mixed effect models to determine which factors of torque pulse application influenced the outcome measures of HE and NPI. As such, we utilized JMP Pro Version 14 (SAS Institute Inc., Cary, NC, USA) to fit a linear mixed model to each of the single-pulse outcome measure data sets consisting of 1280 data points (16 participants x 16 pulse conditions x 5 strides x 1 mean value). In the linear mixed effects models, each mean data point was assigned a unique combination of values from the following effects: stride number (−1, 0, 1, 2, or 3), pulse timing as a phase of gait cycle (Early Stance or Late Stance), knee torque amplitude in N·m (−10, 0, or 10), hip torque amplitude in N·m (−15, 0, or 15), and participant (1 through 16). We added a null data set for hypothetical pulses 17 and 18 in order to implement the linear mixed effect model which requires a full factorial dataset. The hypothetical pulses 17 and 18 consist of zero torque amplitudes in early and late stance, respectively, by averaging measures from pulses 1 through 8 and 9 through 16 from stride -1, respectively. The averaged data from stride -1 was copied to the remaining four strides (0, 1, 2, 3) within each of the 16 participants.

The fixed effects include stride number (Stride), pulse time as phase in gait cycle (Phase), knee torque (Knee), and hip torque (Hip) which therefore provide 4 main effects, 6 twoway effects, 4 three-way effects, and 1 four-way effect. The random effects include the main effect of participant and 4 two-way effects of participant by stride number, pulse time, knee torque, and hip torque. Main and interaction effect terms that are significant at a false positive rate of *α* < 0.05 will be reported.

##### b) Pairwise Tests

To determine if any pulse condition significantly modulated the outcome measure, we performed pairwise tests at the group level between the baseline stride (−1) and each of the following strides (0, 1, 2, 3), at a false positive rate of *α* < 0.05*/*16, given Bonferroni correction for 16 comparisons (one per pulse condition, within a stride condition). The performed pairwise tests were selected on a test by test basis depending upon the normality of the baseline and compared stride data sets. The Shapiro-Wilk parametric hypothesis test of composite normality was utilized to determine the normality of each data set. If both compared data sets were normal, a paired t-tests was performed, otherwise a Wilcoxon signed-rank test was performed.

##### c) Pulse Selection for Repeated-pulse Experiment

Due to concerns on possible participant fatigue and time constraints, only a subset of the original 16 pulse conditions could be tested in the repeated-pulse experimental protocol. As such, we selected a subset of 8 pulse conditions which had modulated the outcome measure in an amplitude-dependent way (positive effect for pulse A, negative effect for pulse A with negative magnitude).

#### 3) Repeated-pulse Experiment

In the repeated-pulse data analysis, the 200 strides of pulse torque intervention is referred to as pulse application, the preceding 100 stride sub-section is baseline, and the 100 stride sub-section following pulse application is after-effects. To perform statistical analysis, we defined five time points of measurement (TP) within each section: baseline (BL) - last 20 strides before intervention, early pulse application (P-E) - strides 2-6 after start of intervention, late pulse application (P-L) - last 5 strides of intervention, early after-effects (AE-E) - strides 2-6 after the end of intervention, and late after-effects (AE-L) - last 5 strides of no pulse condition after intervention. At each of these time points, we obtained the outcome measure as the mean for the designated strides. Stride-averaged outcome measures are used for analysis in Sections II-E3a & II-E3c.

##### a) Linear Mixed Effect Models

The pulses considered in the repeated-pulse experiment do not span factorially all combinations of pulse factors like in the single-pulse experiment. As such, to determine which factors of torque pulse application influenced the primary outcome measures of HE and NPI in the repeated-pulse experiment, we developed two separate linear mixed models, where each model examines the data of four of the eight pulse conditions. Therefore, each model examines a data set of 320 points (16 participants x 4 pulse conditions x 5 time points). The linear mixed effect Model A includes the four pulses where torque is applied to both the hip and knee, at the same time (Table I). Model A effects are pulse time as the phase of gait cycle (Early or Late Stance), knee & hip torque direction (Flex or Ext), time point of measurement (BL, P-E, P-L, AE-E, or AE-L), and participant (1 through 16). The fixed effects include the main, two-way, and three-way effects of pulse time, direction, and time point. The random effects include the main effect of participant and two-way interaction of participant and the main effects. The linear mixed effect Model B includes the four single joint pulse conditions (Table II). Model B effects include knee & hip torque direction, and joint & phase combination (Hip & Early Stance or Knee & Late Stance), time point of measurement, and participant. The fixed effects include the main, two-way, and three-way effects of direction, joint & pulse time combination, and time point of measurement. The random effects include the main effect of participant and twoway interaction between participant and the main effects. Both models for both measures have fixed effects tests conducted with a false positive rate of *α* < 0.05.

**TABLE I:** Model A consisted of the four double-joint pulses with the shown break-down of factors.

| Model A |  | Phase |  |
| --- | --- | --- | --- |
|  |  | Early Stance | Late Stance |
| Direction | Ext | Pulse #5 | Pulse #13 |
|  | Flex | Pulse #8 | Pulse #16 |

**TABLE II:**
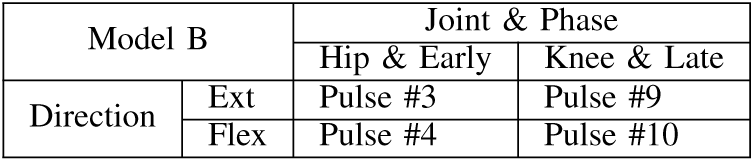
Model B consisted of the four single-joint pulses with the shown break-down of factors.

| Model B |  | Joint & Phase |  |
| --- | --- | --- | --- |
|  |  | Hip & Early | Knee & Late |
| Direction | Ext | Pulse #3 | Pulse #9 |
|  | Flex | Pulse #4 | Pulse #10 |

##### b) Pairwise Tests

Pairwise tests were conducted to establish whether any pulse condition significantly modulated the outcome measures during pulse application (2 paired tests per pulse pairing P-E with BL and P-L with BL), and after pulse application (2 paired tests per pulse pairing AE-E with BL and AE-L with BL). The Shapiro-Wilk test was used to detect normality of the paired samples. If the samples were normal, a t-test was performed, otherwise a Wilcoxon signed-rank test was performed. For either test, a false-positive rate of *α* = 0.05*/*32 was selected, using a Bonferroni correction to account for 32 comparisons (4 comparisons per pulse and 8 pulses).

##### c) Responder Analysis

We sought to establish whether the response of individual participants to the intervention followed a pattern of adaptation or learning (i.e., whether after-effects were in the opposite or in the same direction of effects measured during training). To establish patterns at the individual participant level, we defined the Z-score for phase P-E and AE-E (early pulse application and early after-effects) of each pulse condition for all 16 participants as: 

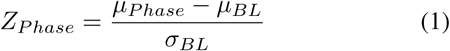

Based on the relative signs of change in outcome measures in the two phases, each participant’s response will follow one of four patterns: positive learning, negative learning, positive adaptation, or negative adaptation (Table III). Each pulse will then have *n* responders for each pattern, with *n* defined as the number of participants whose response to a condition follows a specific pattern. For each pulse, we report which pattern was the one with the largest number of responders, and the mean Z-scores of responders.

**TABLE III:** Definition of response patterns based on Z-scores measured in the early pulse application (P-E), and early after-effects (AE-E).

| Pattern | P-E Z-Score | AE-E Z-Score |
| --- | --- | --- |
| Positive Learning | Positive | Positive |
| Negative Learning | Negative | Negative |
| Positive Adaptation | Negative | Positive |
| Negative Adaptation | Positive | Negative |

## III. Results

### A. Single-pulse Experiment

#### 1) Linear Mixed Effect Models

##### a) HE

The linear mixed effect model for HE had an *R*^2^ adjusted of 0.91. All of the statistically significant fixed effect tests terms are shown in Table S1 for HE. For the significant effects involving stride, the least squares means estimates of levels or post-hoc Tukey HSD pairwise comparisons are presented.

The significant main effect of stride is driven by a reduction in HE during the stride of pulse application (Stride -1: 20.5 ± 1.0 deg, Stride 0: 19.7 ± 1.0 deg, *p* < 0.001). The only significant paired comparisons were measured between HE at stride 0 and all other strides. The significant two-way interaction of stride and knee torque is driven by a significant decrease in HE from stride -1 to 0 for knee flexion torque (change in HE, Knee Flex: -1.8 ± 0.2 deg, *p* < 0.001) which is significantly greater than the decrease in HE for knee extension torque (change in HE, Knee Ext: -0.4 ± 0.2 deg, *p* < 0.001). The significant two-way interaction of stride and hip torque is driven by a significant decrease in HE from stride -1 to 0 for hip flexion torque (change in HE, Hip Flex: -2.2 ± 0.2 deg, *p* < 0.001) which is significantly more negative than the change in HE from stride -1 to 0 for hip extension torque (change in HE, Hip Ext: 0.2 ± 0.2 deg, *p* < 0.001). The significant two-way interaction of stride and phase is driven by a significant decrease in HE from stride -1 to 0 in early stance (change in HE: Early Stance: -1.1 ± 0.2 deg, *p* < 0.001).

The significant three-way interaction of stride, phase, and knee torque is driven by a significant decrease in HE from stride -1 to 0 for knee extension torque at early stance (change in HE: Early Stance, Knee Ext: -2.3 ± 0.3 deg, *p* < 0.001), by a significant increase in HE for knee extension torque at late stance (change in HE, Late Stance, Knee Ext: 1.6 ± 0.3 deg, *p* < 0.001), and by a significant decrease in HE for knee flexion torque at late stance (change in HE, Late Stance, Knee Flex: -2.9 ± 0.3 deg, *p* < 0.001). The significant three-way interaction of stride, phase, and hip torque is driven by a significant decrease in HE from stride -1 to 0 for hip extension torque at early stance (change in HE: Early stance, Hip Ext: -2.4 ± 0.3 deg, *p* < 0.001), by a significant increase in HE for hip extension at late stance (change in HE: Late stance, Hip Ext: 2.8 ± 0.3 deg, *p* < 0.001), and by a significant decrease in HE for hip flexion at late stance (change in HE: Late stance, Hip Flex: -4.1 ± 0.3 deg, *p* < 0.001).

##### b) NPI

The linear mixed effect model for NPI had an *R*^2^ adjusted of 0.98. All of the statistically significant fixed effect terms are shown in Table S2 for NPI. For the significant effects involving stride, the least squares means estimates of levels or post-hoc Tukey HSD pairwise comparisons are presented.

The significant main effect of stride is driven by an increase in NPI during the stride of pulse application (Stride -1: 28.7 ± 2.1 ms, Stride 0: 29.5 ± 2.1 ms, *p* < 0.001). The significant paired comparisons were measured between NPI at stride 0 and all other strides. The significant two-way effect of stride and knee torque is driven by a significant increase in NPI from stride -1 to 0 for knee flexion torque (change in NPI: Knee Flex: 1.8 ± 0.2 ms, *p* < 0.001) which is significantly greater than the change measured for knee extension torque from stride -1 to 0 (change in NPI: Knee Ext: 0.28 ± 0.20 ms, *p* < 0.001). The significant two-way interaction of stride and hip torque is driven by a significant increase in NPI from stride -1 to 0 in hip extension torque (change in NPI: Hip Ext: 1.7 ± 0.2 ms, *p* < 0.001) which is significantly greater than the change measured for hip flexion torque (change in NPI: Hip Flex 0.1 ± 0.2 ms, *p* < 0.001). The significant two-way interaction of stride and phase is driven by a significant increase in NPI from stride -1 to 0 in early stance (change in NPI: Early Stance: 1.1 ± 0.2 ms, *p* < 0.001) which is a significantly greater than the increase measured in late stance (change in NPI: Late Stance: 0.5 ± 0.2 ms, *p* = 0.008).

The significant three-way interaction of stride, phase, and knee torque is driven by a significant increase in NPI from stride -1 to 0 for early stance knee extension torque (change in NPI: Early Stance, Knee Ext: 3.4 ± 0.3 ms, *p* < 0.001), by a significant decrease in NPI for late stance knee extension torque (change in NPI: Late Stance, Knee Ext: -2.9 ± 0.3 ms, *p* < 0.001) and by a significant increase in NPI for late stance knee flexion torque (change in NPI: Late Stance, Knee Flex: 4.2 ± 0.3 ms, *p* < 0.001). The significant three-way interaction of stride, phase, and hip torque is driven by a significant increase in NPI from stride -1 to 0 for early stance hip extension torque (change in NPI: Early Stance, Hip Ext: 2.8±0.3 ms, *p* < 0.001).

#### 2) Pairwise Tests

The outcome measure of HE had statistically significant pairwise comparisons present in seven of the sixteen total pulse conditions (Fig. 5). The only significant pairwise differences were between baseline stride (−1) and the stride of pulse application (0), as seen in Table IV. Pulse conditions 1 and 5, both containing knee extension torque pulses in early stance, decreased HE. Flexion pulses at late stance, conditions 10 and 12, both decreased HE, and combine to form pulse condition 16 which had an even greater decrease in HE. Pulse conditions 11 and 13 both feature late stance hip extension torque pulses and increased HE.

**TABLE IV:** The expected effect on TLA based on our previous work [10], effect size, and *p* value for the pulse of application relative to baseline for measured HE and NPI, (threshold *p* = 0.05*/*16 = 0.003) for each of the 16 pulse conditions in single-pulse application

| Pulse # | Expected effect on TLA | HE Effect Size | HE p Value | NPI Effect Size | NPI p Value |
| --- | --- | --- | --- | --- | --- |
| 1 | Increase | -0.999 | 0.001 | 1.198 | <0.001 |
| 2 | Decrease | -0.184 | 0.472 | -0.297 | 0.379 |
| 3 | Increase | -0.540 | 0.047 | 1.852 | <0.001 |
| 4 | Decrease | 0.095 | 0.709 | -0.692 | 0.014 |
| 5 | Increase | -1.599 | <0.001 | 1.413 | <0.001 |
| 6 | Unknown | -0.436 | 0.101 | 0.786 | 0.010 |
| 7 | Unknown | -0.487 | 0.070 | 0.927 | 0.003 |
| 8 | Decrease | -0.057 | 0.823 | -0.629 | 0.023 |
| 9 | Decrease | 0.583 | 0.034 | -1.836 | <0.001 |
| 10 | Increase | -1.317 | <0.001 | 3.530 | <0.001 |
| 11 | Increase | 1.781 | <0.001 | 0.065 | 0.836 |
| 12 | Decrease | -1.487 | <0.001 | 0.037 | 0.679 |
| 13 | Unknown | 1.597 | <0.001 | -1.190 | <0.001 |
| 14 | Decrease | -0.508 | 0.020 | -2.193 | <0.001 |
| 15 | Increase | 0.081 | 0.749 | 3.392 | <0.001 |
| 16 | Unknown | -1.622 | <0.001 | 2.081 | <0.001 |

**Fig. 3:**
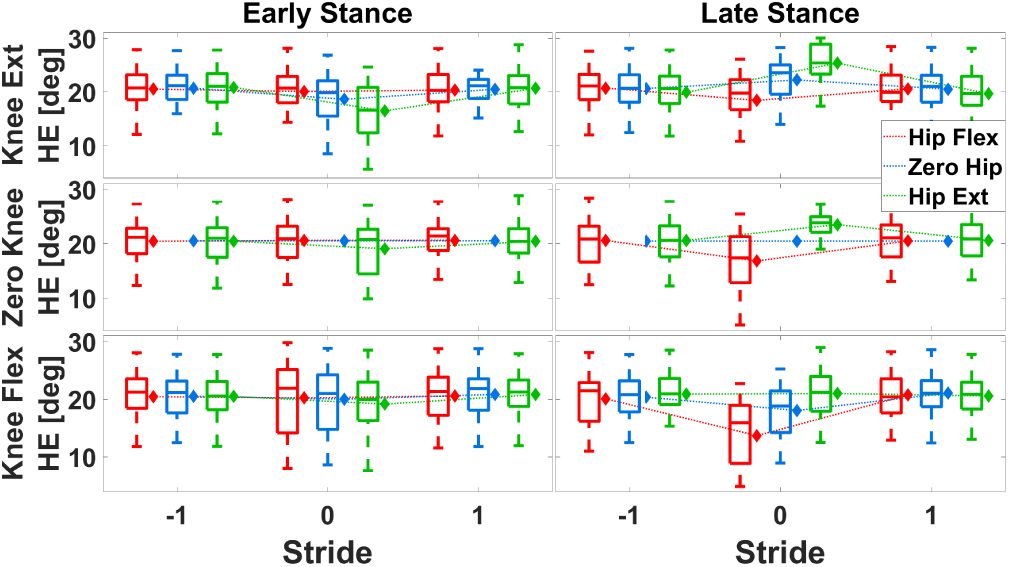
Breakdown of HE measurements (boxplots) and model estimates (diamonds) for different levels of the four factors considered in the single-pulse experiment.

**Fig. 4:**
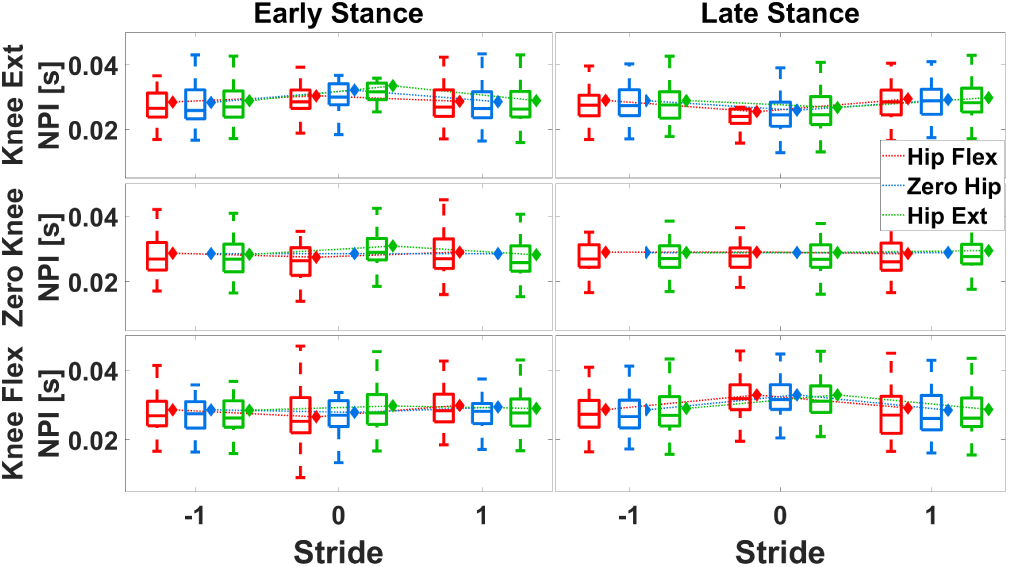
Breakdown of NPI measurements (boxplots) and model estimates (diamonds) for different levels of the four factors considered in the single-pulse experiment.

**Fig. 5:**
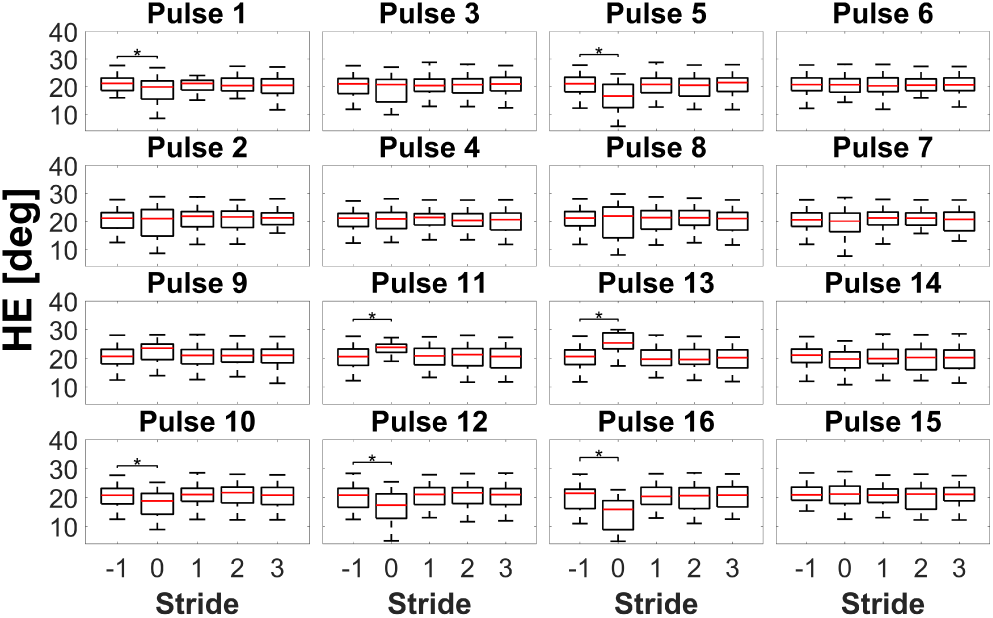
The group-wise depiction of HE by stride for all pulse conditions in the single-pulse experiment. Asterisks indicate a statistically significant comparison between the baseline stride and specified following stride.

The outcome measure of NPI had statistically significant pairwise comparisons in seven pulse conditions, only between baseline stride (−1) and the stride of pulse application (0) (Fig. 6). Hip extension at early stance lead to a significant increase in NPI. Pulse conditions 9, 13, and 14 all contained late stance knee extension torque and lead to a decrease in NPI. Conversely, pulse conditions 10, 15, and 16 all contain late stance knee flexion torque and lead to an increase in NPI.

**Fig. 6:**
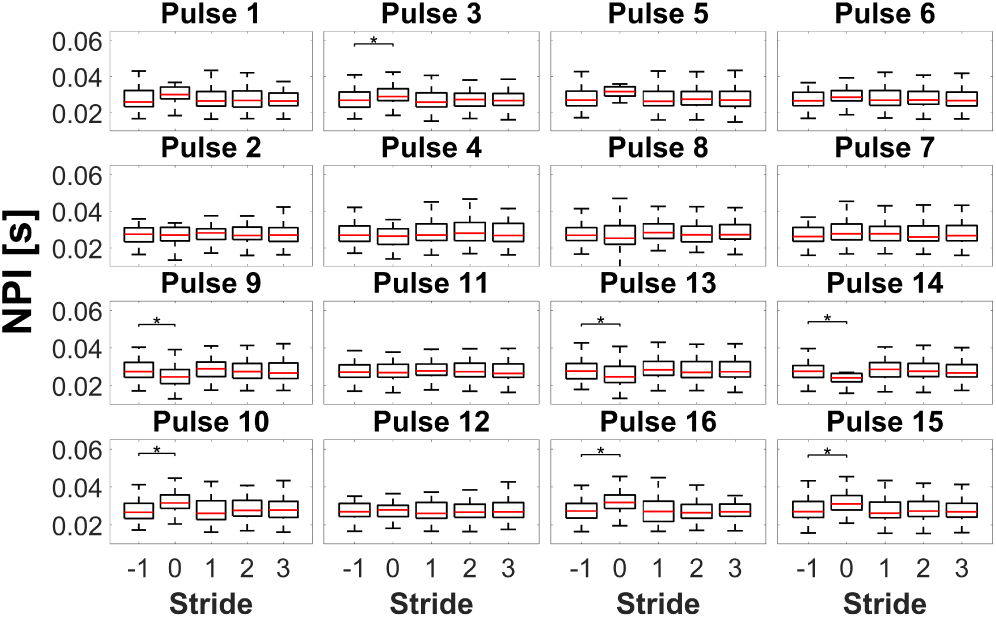
The group-wise depiction of NPI by stride for all pulse conditions in the single-pulse experiment. Asterisks indicate a statistically significant comparison between the baseline stride and specified following stride.

#### 3) Pulse Selection for Repeated-pulse Experiment

The selected subset of 8 pulse conditions include four pulse conditions with a positive modulation in NPI and negative modulation in HE: 3, 5, 10, and 16. The other four pulse conditions had a negative modulation in NPI and positive modulation HE: 4, 8, 9, and 13. These selected pulse conditions are depicted in Fig. 2 where the pulse title colors are red or green for negative and positive modulation, respectively, of NPI in single pulse application.

### B. Repeated-pulse Experiment

The group means by stride for HE and NPI are shown for all eight repeated-pulse conditions in Figs. 7 & 9.

**Fig. 7:**
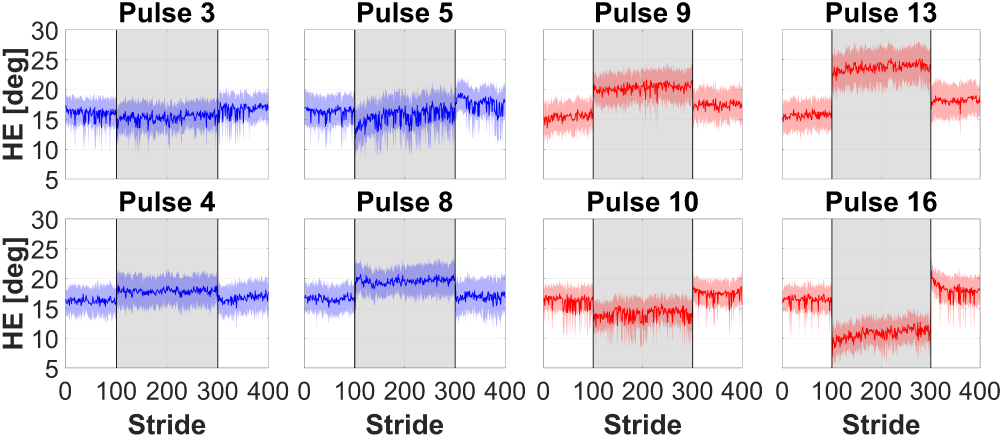
Group mean HE data by stride for all eight repeated-pulse conditions.

#### 1) Linear Mixed Effect Model

The linear mixed effect Models A & B for HE have an *R*^2^ adjusted of 0.81 and 0.68, respectively. The linear mixed effect Models A & B for NPI have an *R*^2^ adjusted of 0.66 and 0.30, respectively. The statistically significant fixed effects tests for all four models are given in Table S3. The linear mixed model estimates of means with standard errors are given in Figs. 8 & 10

**Fig. 8:**
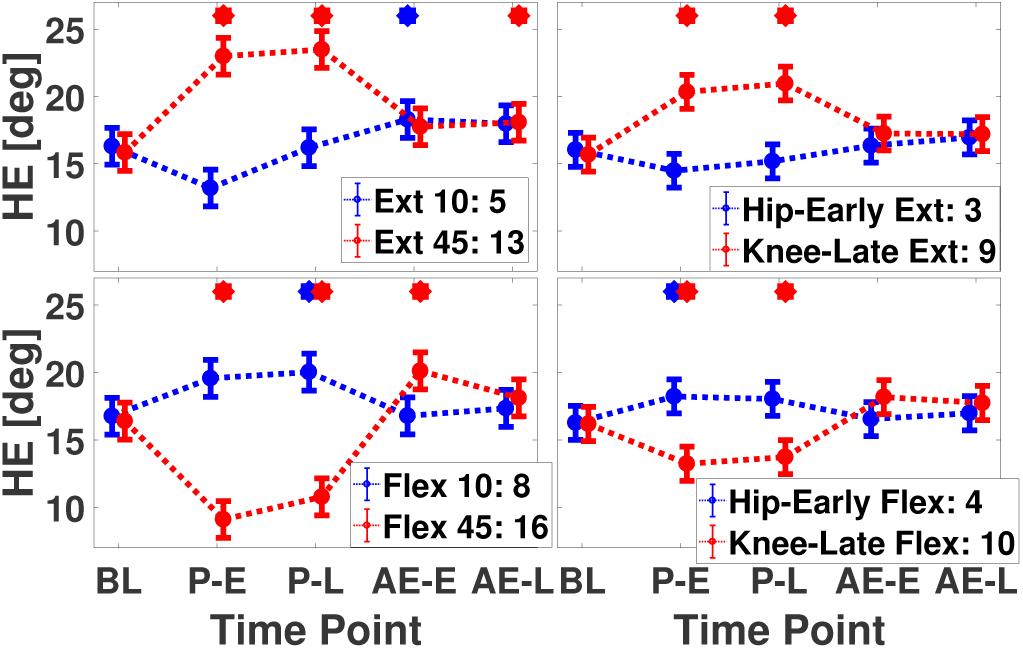
Breakdown of HE by factor for the eight pulses tested in the repeated-pulse experiment. Diamonds indicate model-estimated means, whiskers indicate s.e.m., asterisks indicate statistically significant pairwise comparison to respective base-line estimate.

The significant two-way interaction in Model A for HE of time point of measurement and direction is driven by multiple pairwise comparisons within direction and between BL and post-BL time points. For extension pulses, HE significantly increased at all time points, compared to baseline: P-E (2.0± 0.6 deg, *p* = 0.030), P-L (3.8 ±0.6 deg, *p* < 0.001), AE-E (2.0± 0.6 deg, *p* = 0.042), AE-L (2.0± 0.6 deg, *p* = 0.041). For flexion pulses, the only significant pairwise comparison is a decrease in HE relative to BL at P-E (−2.2 ± 0.6 deg, *p* = 0.009). The significant three-way interaction in Model A of time point of measurement, phase, and direction is driven by six significant pairwise comparisons. For extension at late stance, HE increased significantly relative to BL at P-E (7.2 ±0.9 deg, *p* < 0.001) and P-L (7.7 ±0.9 deg, *p* < 0.001). Within the flexion direction and early stance phase, HE increased significantly relative to BL at P-L (3.3±0.9 deg, *p* = 0.044). Within the flexion direction and late stance phase, HE significantly changed relative to BL at P-E (−7.3 ± 0.9 deg, *p* < 0.001), P-L (−5.6 ± 0.9 deg, *p* < 0.001), and AE-E (3.7 ± 0.9 deg, *p* = 0.008).

The main effect in Model B for HE of time point of measurement is significant in which P-L (17.0 ± 1.1 deg, *p* = 0.017), AE-E (17.1 ± 1.1 deg, *p* = 0.006), and AE-L (17.2 ± 1.1 deg, *p* = 0.002) all are of significantly greater HE than BL (16.0 ± 1.1 deg). The two-way effect in Model B of time point of measurement and direction pairwise comparison is an increase in HE within extension, from BL to P-L (2.2 ± 0.5 deg, *p* = 0.003). The three-way interaction between time point of measurement, joint & phase, and direction is driven by an increase of HE measured for knee & late stance extension pulses, relative to BL, during P-E (4.7±0.9 deg *p* < 0.001) and P-L (5.3±0.9 deg *p* < 0.001).

In Model A for NPI, there are no significant between time point of measurement and within phase and direction pairwise comparisons to report. Similarly, in Model B for NPI, there are no significant between time point of measurement and within joint & phase and direction pairwise comparisons to report.

#### 2) Pairwise Tests

The effect size of the change in HE and NPI between BL and post-BL phases are reported in Table V. Five out of eight pulses significantly modulated HE at P-E, with three pulses (4, 9, and 13 - all including early stance flexion or late stance extension) inducing a positive change, and two pulses (10 and 16, all including late stance flexion) inducing a negative change. The effects of these pulses were sustained at P-L for all pulses except pulse 4. Pulse 8 (early stance flexion) exhibited a positive modulation in HE at P-L following a non-significant, but of similar magnitude, positive modulation at P-E. Significant changes in HE were measured at AE-E for two pulses - 5 (positive increase in HE, in the opposite direction of non-significant effects measured during P conditions), and 16 (positive increase in HE, in the opposite direction of effects measured during P conditions). One pulse (13) showed a significant change in HE in AE-L, resulting from a positive increase in HE, in the same direction of effects measured during P conditions.

**TABLE V:** The effect size for all pairwise comparisons between baseline and each following phase for all eight repeated-pulse conditions. Values are bolded if statistically significant at *p* = 0.05/32.

| Pulse # | HE |  |  |  | NPI |  |  |  |
| --- | --- | --- | --- | --- | --- | --- | --- | --- |
|  | P-E | P-L | AE-E | AE-L | P-E | P-L | AE-E | AE-L |
| 3 | -0.774 | -0.288 | 0.098 | 0.607 | 0.419 | 0.247 | 0.007 | 0.677 |
| 4 | **1.484** | 0.889 | 0.117 | 0.429 | -0.377 | -0.141 | 0.728 | 0.204 |
| 5 | -0.815 | -0.045 | **1.562** | 0.912 | 0.213 | 0.020 | -0.087 | 0.230 |
| 8 | 0.944 | **1.021** | 0.007 | 0.208 | -0.195 | 0.537 | 0.816 | 1.080 |
| 9 | **2.326** | **1.721** | 0.815 | 0.762 | -0.382 | -0.154 | 0.340 | 0.243 |
| 10 | **-1.729** | **-1.057** | 0.806 | 0.954 | **1.561** | 0.310 | -0.068 | 0.163 |
| 13 | **2.272** | **1.960** | 0.889 | **1.569** | -0.447 | 0.018 | 0.154 | 0.294 |
| 16 | **-2.553** | **-1.908** | **1.333** | 0.674 | **1.566** | 0.209 | -0.277 | 0.419 |

Two out of the eight pulses significantly modulated NPI at P-E, with both pulses (10 and 16 - both including flexion) inducing a positive change.

#### 3) Responder Analysis

For the measure of HE, pulse conditions 3, 5, 10 and 16 have the dominant pattern of positive adaptation, with high proportions of the participants following the pattern: 12, 12, 14, and 14 participants, respectively. The other four pulse conditions (4, 8, 9, and 13) have the dominant pattern of positive learning, with 9, 9, 12, and 11 participants following the pattern, respectively (Fig. 11 & Table VI).

**TABLE VI:**
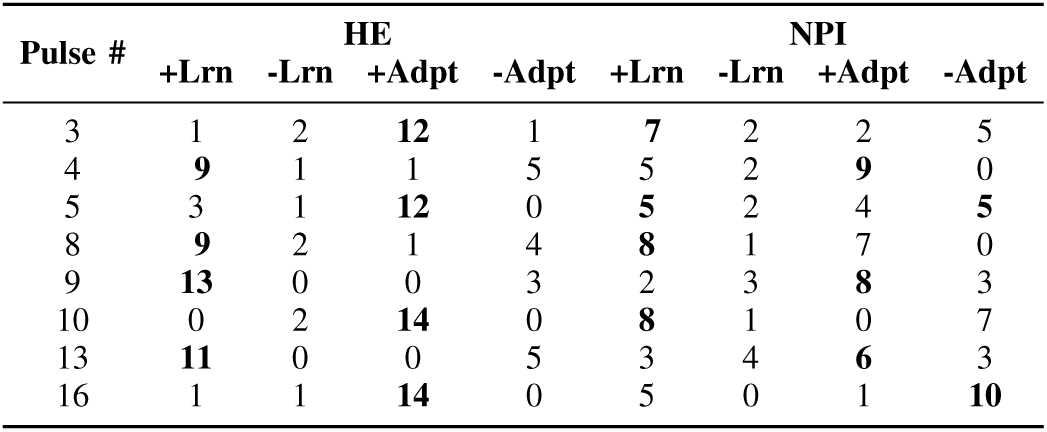
Number of repeated-pulse responders for each behavioral pattern for each pulse condition and outcome measure. Bold entries indicate the dominant pattern for a specific pulse condition and outcome measure.

**Fig. 9:**
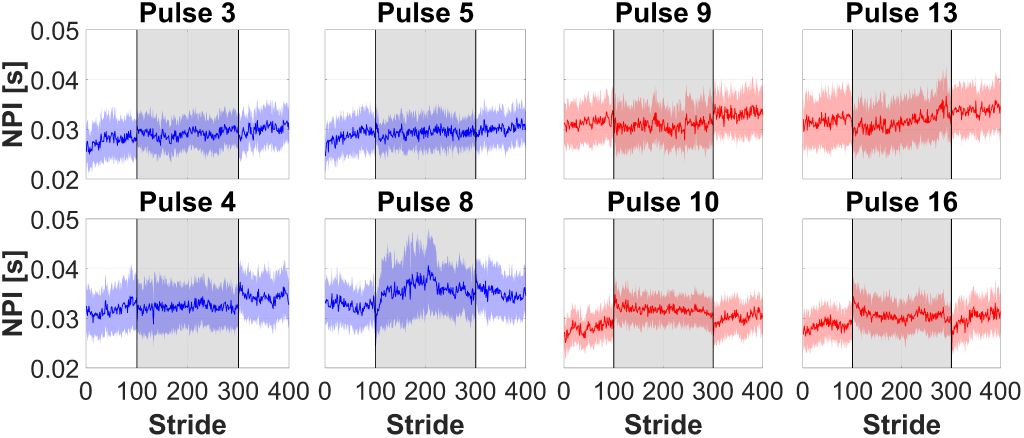
Group mean NPI data by stride for all eight repeated-pulse conditions.

**Fig. 10:**
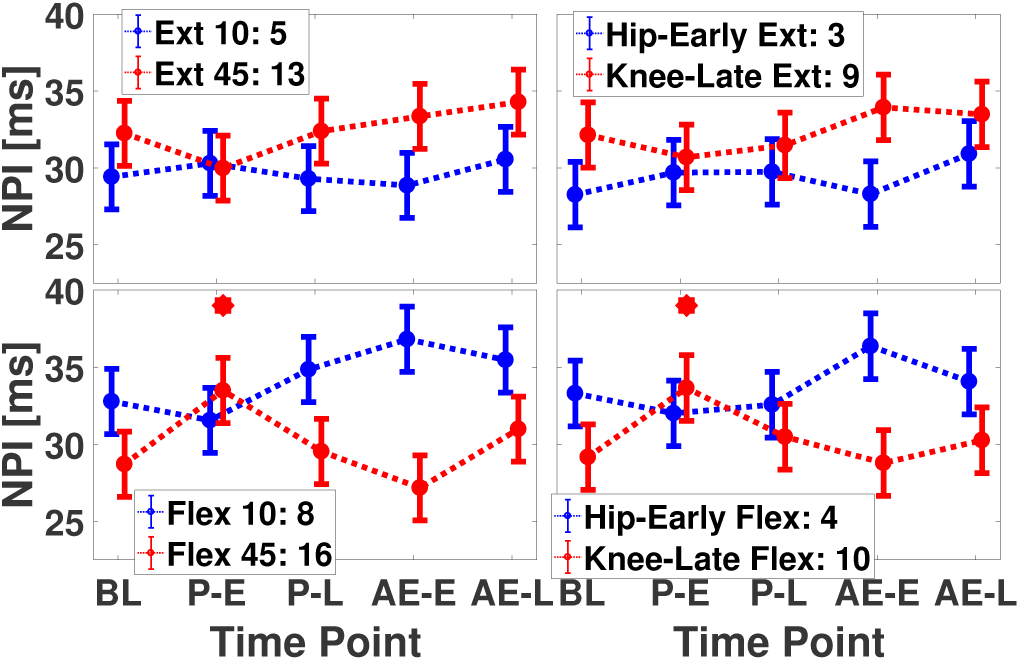
Breakdown of NPI by factor for the eight pulses tested in the repeated-pulse experiment. Diamonds indicate model-estimated means, whiskers indicate s.e.m., asterisks indicate statistically significant pairwise comparison to respective base-line estimate.

**Fig. 11:**
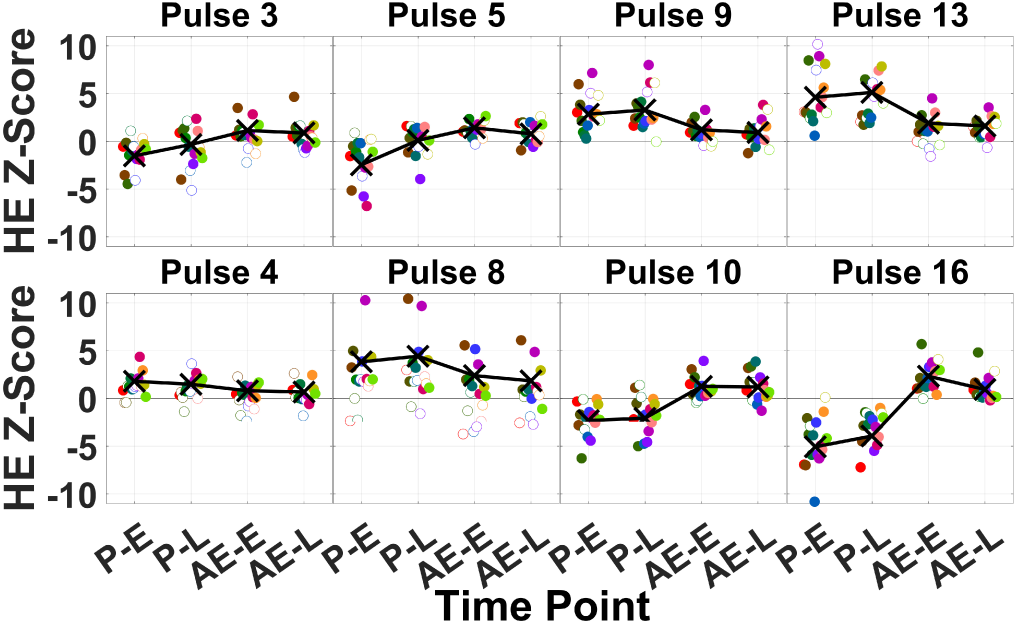
Z-scores of responders to the dominant pattern for HE. Filled circles are individual subject *z*-scores, crosses are group mean Z-scores.

For the measure of NPI, pulse conditions 3, 8, and 10 all have the dominant pattern of positive learning with 7, 8, and 8 participants following the pattern, respectively. Pulse conditions 4, 9, and 13 all have the dominant pattern of positive adaptation with 9, 8, and 6 participants following the pattern, respectively. The dominant pattern of pulse condition 16 is negative adaptation with a total of 10 participants following the pattern. As for pulse condition 5, there was an equal number of participants following the two dominant patterns of positive learning and negative adaptation with a total of 5 participants for each (Fig. 12 & Tables VI)

**Fig. 12:**
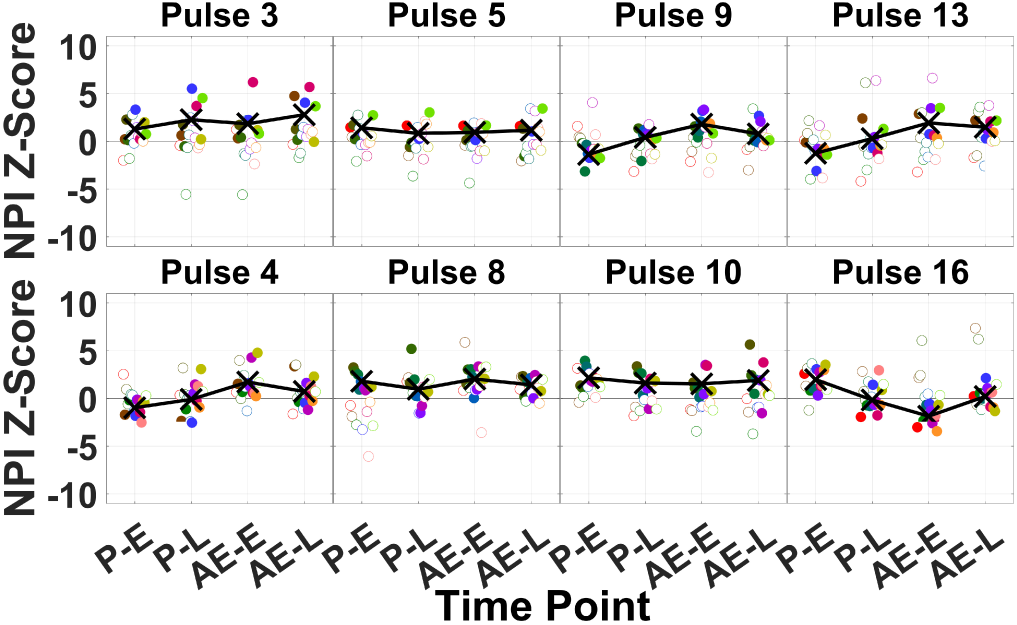
Z-scores of responders to the dominant pattern for NPI. Filled circles are individual subject *z*-scores, crosses are group mean Z-scores.

## IV. Discussion and conclusion

In this work, we used a robotic exoskeleton to apply torque pulses to the hip and knee joint during stance with the intention of modulating propulsion dynamics. The selection of the set of pulses to test was informed by our prior work, where we investigated the differences in joint moments associated with modulation of push-off posture (a biomechanical component of propulsion) during walking. We conducted two experiments to measure the effects of torque pulses on biomechanical measures of propulsion, defined by normalized propulsive impulse (NPI), and hip extension (HE). In the first experiment (single-pulse experiment), we applied pulses only during single strides to quantify the intrinsic effects of the intervention - i.e., the effects measured under minimal changes in the subject’s neuromotor coordination. In the second experiment (repeated-pulse experiment), we applied pulses continuously over multiple strides, to measure how subjects would adapt to robotic training, and how they would respond right after exposure to the training. Effects measured during the stride of pulse application were mostly in the same direction across the two experiments, while the effects measured after exposure to torque pulses largely differed across the two experiments. We will discuss the effects during pulse application and after pulse application, for both experiments, separately, in the following sections.

### A. Effects during pulse application

Several features of the effects of pulse conditions on NPI and HE were qualitatively similar across the two experiments during pulse application. In fact, the changes in both outcome measures for the single-pulse experiment were in the same direction as the changes measured in the first few strides during the repeated-pulse experiment for all conditions except one, pulse 8 - HE, whose effect size in the single-pulse experiment was close to zero (−0.057). Interestingly, the effects measured in NPI aligned with the expected modulation of TLA based on our previous biomechanical investigation [9], [10], while the effects measured in HE were usually in the opposite direction as NPI. The changes in HE and NPI measured in the repeated-pulse experiment aligned with the single-pulse experiment during pulse application, and included both positive and negative effects. Instead, after-effects measured in the repeated-pulse experiment were primarily associated to positive learning or positive adaptation, with no significant negative changes in any outcome measure.

As per which features of the applied pulses modulate propulsion, we found that timing of pulse application (early or late stance) was a significant factor, as a given condition at early stance often induced an opposite effect on the outcome measures when applied at late stance. Consistently in both experiments, extension torque applied to the hip and/or knee joint at late stance increased HE during pulse application — range of change (RoC) in HE: 1.6 ± 0.3 deg to 7.7 ± 0. deg or 7.8 ± 1.5% to 47.5 ± 5.6%, *p* < 0.001. In a partially mirrored fashion, extension torque applied to the hip and/or knee joint at early stance resulted in a decreased HE during pulse application. The direction of the effects was consistent between single- and repeated-pulse experiments, but post-hoc comparisons were statistically significant only in the single-pulse experiment — RoC HE: (−2.3, −2.4) ± 0.3 deg or (−11.2, −11.7) ± 1.5%, *p* < 0.001 for all conditions involving extension torque at early stance.

Effects of flexion torque pulses, and effects of all conditions on NPI were not as uniform across joint and between the two experiments, so more granular effects will be discussed separately below. In the single-pulse experiment, the effects of extension torque pulses on NPI were in the opposite directions as the effects of the same conditions on HE, with a significant decrease in NPI during pulse application for pulses applied at late stance (change in NPI: −2.9 ± 0.3 ms or −10.0 ± 0.9%, *p* < 0.001 for knee torque, no significant change for hip torque), and a significant increase in NPI for pulses applied at early stance — RoC in NPI: (2.8, 3.4) ± 0.3 ms or (9.8, 12.0) ±0.9%, *p* < 0.001 for both knee and hip torque). These conditions, however, did not translate in a significant change in NPI during pulse application for any extension condition in the repeated-pulse experiment.

As per the effects of flexion pulses, in the single-pulse experiment, only knee flexion torque pulses at late stance significantly decreased HE (change in HE: 2.9 ± 0.3 deg or 10.1 ± 0.9%, *p* < 0.001). This result was confirmed in the repeated stride experiment — RoC HE: (−5.6, −7.3) ± 0.9 deg or (−34.6, −45.0) ±5.6%, *p* < 0.001 for both early and late conditions) — in addition to a significant increase in NPI resulting from flexion pulses applied at late stance (RoC in NPI: (4.5, 4.8) ±2.0 ms or (14.6, 15.6) ±6.5%, *p* < 0.001 for both pulse conditions). Opposite effects were measured on NPI for the same conditions, with knee flexion torque pulses at late stance significantly increasing NPI (change in NPI: 4.2 ± 0.3 ms or 14.5 ± 0.9%, *p* < 0.001 in the single-pulse experiment), paralleled by the increase in NPI measured in torque pulse conditions including flexion at late stance (16 and 10) in the repeated pulses experiment.

Small and non statistically significant effects in HE and NPI were measured for flexion pulses applied at early stance in the single-pulse experiment. Instead, post-hoc comparisons involving flexion pulses at early stance showed significant increase in HE for hip flexion torque (change in HE: 3.3 ± 0.9 deg or 20.4 5. ± 6%, *p* = 0.044) in the repeated-pulse experiment.

A possible explanation for the alignment between the measured modulation of NPI and the expected modulation of TLA could be the fact that the applied torque pulses effectively modulated TLA, thus facilitating the generation of increased NPI with a roughly constant net ankle moment. The fact that the measured effect on HE was on the opposite direction as expected may arise from the fact that HE is only a surrogate for TLA, confounded by variables such as knee and ankle angles that may be crucially modulated during push-off.

The agreement of the effects in HE and NPI between the two experiment suggests that effects during pulse application, especially in the first strides, are primarily due to biomechanics, and influenced by the subject response only to a small extent. Certainly, the active response of subjects plays a larger role during later strides of pulse application, where some NPI effects cease to be significant. The decrease of significance of effects in NPI in later strides is certainly also influenced by physical constraints of the experimental setup. In fact, the imposed treadmill belt speed, combined with the limited anterior lunge of the robot, could lead the participant to reduce propulsive effort such that they cease to infringe upon the motion constraints of the setup.

### B. Effects after pulse application

In the single-pulse experiment, we did not measure any significant modulation in outcome measures in the strides following pulse application (Fig. 5 and Fig. 6). Instead, in the repeated-pulse experiment, we measured significant changes in outcome measures in three of the eight pulses applied, all for HE. Significant after-effects in HE were only measured in the positive direction, as a result of the application of hip and knee extension torque in early or late stance, and as the result of the application of hip and knee flexion torque at late stance. Interestingly, one only condition (hip and knee extension torque at late stance), yielded after-effects in HE that persisted after the first few strides of training. Interestingly, this was the experimental condition that yielded the largest positive change in HE during pulse application both in the single-pulse and in the repeated-pulse experiment. While no pulse torque condition modulated NPI after pulse application significantly at the group level, a responder analysis confirmed that the dominant pattern of individual subject response was consistent with a positive change in NPI after pulse application, with six pulses inducing a response classified as “positive learning” or “positive adaptation”, with only one pulse inducing primarily a response of “negative adaptation”.

Of the four pulses that induced a response in HE classified as “positive learning”, three of them (knee extension at late stance, hip and knee extension at late stance, and hip flexion at late stance) induced a response in NPI classified as “positive adaptation”, while one (hip and knee flexion at early stance) induced a response in NPI classified as “positive learning”. As such, while only one of these pulses produced statistically significant changes in HE at the group level, all four pulses that induced positive learning on HE resulted in positive change in NPI for individual subjects.

Of the four pulses that induced a response in HE classified as “positive adaptation”, two of them (hip extension at early stance, knee flexion at late stance) induced a response in NPI classified as “positive learning”. As such, while none of these pulses produced statistically significant changes in HE at the group level, two of these pulses resulted in positive change in NPI for individual subjects. Pulse 16 (hip and knee flexion at late stance) was the only condition that produced a dominant response featuring negative after-effects (10/16 subjects).

Overall, the effects measured after pulse application are mostly in the direction of a positive change in both HE and NPI. The magnitude of the effects observed in HE are greater than those measured in NPI, resulting in no pulse condition with a statistically significant change in NPI at the group level. However, it is possible that the magnitude of the effects in NPI are reduced due to the fact that this experiment was conducted at fixed speed. In a fixed speed experiment, even if the subject has been trained to modulate NPI in a positive direction, maintaining an increased NPI would result in the subject walking anteriorly toward the limit of the treadmill. In our experimental setup, also, subjects are further constrained by the limited range of motion of the anterior lunge DOF of the ALEX II robot. Both constraints make it unlikely to be able to observe sustained effects in NPI in experiments conducted on a fixed speed treadmill. As such, we wish to form a future protocol to promote successful NPI modulation through the implementation of an adaptive treadmill controller, which has the capacity to increase the treadmill belt speed in direct response to the real-time gait parameters of the participant [14].

### C. Study limitations

This study had limitations which could be addressed through protocol modifications in future work. First, the statistical model describing effects of single joint pulses during the repeated-pulse experiment on NPI was able to account only for a limited portion of the total variance in the dataset (*R*^2^ adjusted = 0.3). This is likely due to the NPI measure being quite noisy, and to the small effect of single joint pulses on NPI, which yields a low signal-to-noise ratio. Second, while it is know that push-off posture of the trailing limb is associated with propulsion, we were unable to directly measure TLA due to limitations in the experimental setup. As such, we utilized the surrogate measure of HE as measured by the robot. Lastly, the set of pulses tested in the repeated-pulse experiment did not include all of the pulses tested in the single-pulse experiment. This choice was done in light of practical limitations in experimental protocol duration and subject availability, and pulses were selected based on the results of the single-pulse experiment. However, as a consequence of this necessary choice, some pulse conditions with potentially strong effects on HE and NPI, such as those combining hip extension and knee flexion torque (pulses 14 and 15 for late stance, and 6 and 7 for early stance). Moreover, the lack of a full factorial set of conditions in the repeated-pulse experiment prevented us from being able to test all interactions between factors, as done for the single-pulse experiment.

## Acknowledgment

This work was supported by the National Science Foundation (NSF-NRI-1638007) and by an Institutional Development Award (IDeA) from the National Institute of General Medical Sciences of the National Institutes of Health (NIH-U54-GM104941).

## V. Supplementary materials

### A. Single-Pulse Fixed Effects

**TABLE S1:** Significant fixed effects estimated via the linear mixed effect model for HE in the single-pulse experiment.

| HE Fixed Effect Tests | Nparm | DFNum | DFDen | F Ratio | Prob >F |
| --- | --- | --- | --- | --- | --- |
| Hip | 2 | 2 | 30 | 15.347 | <0.0001 |
| Stride | 4 | 4 | 60 | 7.191 | <0.0001 |
| Stride-Knee | 8 | 8 | 1200 | 10.593 | <0.0001 |
| Stride-Hip | 8 | 8 | 1200 | 16.851 | <0.0001 |
| Phase-Knee | 2 | 2 | 1200 | 17.062 | <0.0001 |
| Phase-Hip | 2 | 2 | 1200 | 49.042 | <0.0001 |
| Stride-Phase | 4 | 4 | 1200 | 3.467 | 0.0080 |
| Stride-Phase-Knee | 8 | 8 | 1200 | 24.896 | <0.0001 |
| Stride-Phase-Hip | 8 | 8 | 1200 | 56.171 | <0.0001 |

**TABLE S2:** Significant fixed effects estimated via the linear mixed effect model for NPI in the single-pulse experiment.

| NPI Fixed Effect Tests | Nparm | DFNum | DFDen | F Ratio | Prob >F |
| --- | --- | --- | --- | --- | --- |
| Hip | 2 | 2 | 30 | 8.090 | 0.0015 |
| Knee | 2 | 2 | 30 | 3.785 | 0.0342 |
| Stride | 4 | 4 | 60 | 8.841 | <0.0001 |
| Stride-Knee | 8 | 8 | 1200 | 11.138 | <0.0001 |
| Stride-Hip | 8 | 8 | 1200 | 11.716 | <0.0001 |
| Phase-Knee | 2 | 2 | 1200 | 67.976 | <0.0001 |
| Stride-Phase | 4 | 4 | 1200 | 4.041 | 0.0029 |
| Stride-Phase-Knee | 8 | 8 | 1200 | 114.255 | <0.0001 |
| Stride-Phase-Hip | 8 | 8 | 1200 | 9.382 | <0.0001 |

### B. Repeated-Pulse Fixed Effects

**TABLE S3:** The fixed effect tests results for the repeated-pulse linear mixed effect models: HE Model A, HE Model B, NPI Model A, and NPI Model B.

| HE-A Fixed Effects Tests | Nparm | DFNum | DFDen | F Ratio | Prob >F |
| --- | --- | --- | --- | --- | --- |
| Direction | 1 | 1 | 15 | 8.938 | 0.009 |
| Phase | 4 | 4 | 60 | 12.730 | <0.001 |
| TP·Direction | 4 | 4 | 195 | 12.751 | <0.001 |
| Phase·Direction | 1 | 1 | 195 | 115.198 | <0.001 |
| TP·Phase·Direction | 4 | 4 | 195 | 68.615 | <0.001 |
| HE-B Fixed Effects Tests | Nparm | DFNum | DFDen | F Ratio | Prob >F |
| Direction | 1 | 1 | 15 | 6.243 | 0.025 |
| TP | 4 | 4 | 60 | 5.338 | 0.001 |
| TP·Direction | 4 | 4 | 195 | 3.934 | 0.004 |
| Joint&Phase·Direction | 1 | 1 | 195 | 43.957 | <0.001 |
| TP·Joint&Phase·Direction | 4 | 4 | 195 | 21.097 | <0.001 |
| NPI-A Fixed Effects Tests | Nparm | DFNum | DFDen | F Ratio | Prob >F |
| Direction | 1 | 1 | 15 | 5.416 | 0.034 |
| Phase·Direction | 1 | 1 | 195 | 38.630 | <0.001 |
| TP·Phase·Direction | 4 | 4 | 195 | 5.392 | <0.001 |
| NPI-B Fixed Effects Tests | Nparm | DFNum | DFDen | F Ratio | Prob >F |
| Direction | 1 | 1 | 15 | 6.311 | 0.024 |
| Joint&Phase·Direction | 1 | 1 | 195 | 20.258 | <0.001 |
| TP·Joint&Phase·Direction | 4 | 4 | 195 | 2.813 | 0.027 |

### C. Validation of Torque Pulse Application

The physical application of torque by the robot was observed to be delayed by approximately 100ms behind the torque command sent to the low-level controller. In order to compensate for this delay, the commanded torque profile preceded the desired torque profile by 10% GC in the single-pulse protocol and 100ms in the repeated-pulse protocol. Shown in Figs. S1 & S2 are the joint torque profiles of the no pulse condition and two commanded demonstration pulse conditions for the knee and hip of a sample participant during the single-pulse protocol. All depicted data are mean curves with standard deviation shading for a compiled set of ten sample single-pulse protocol gait cycles.

**Fig. S1:**
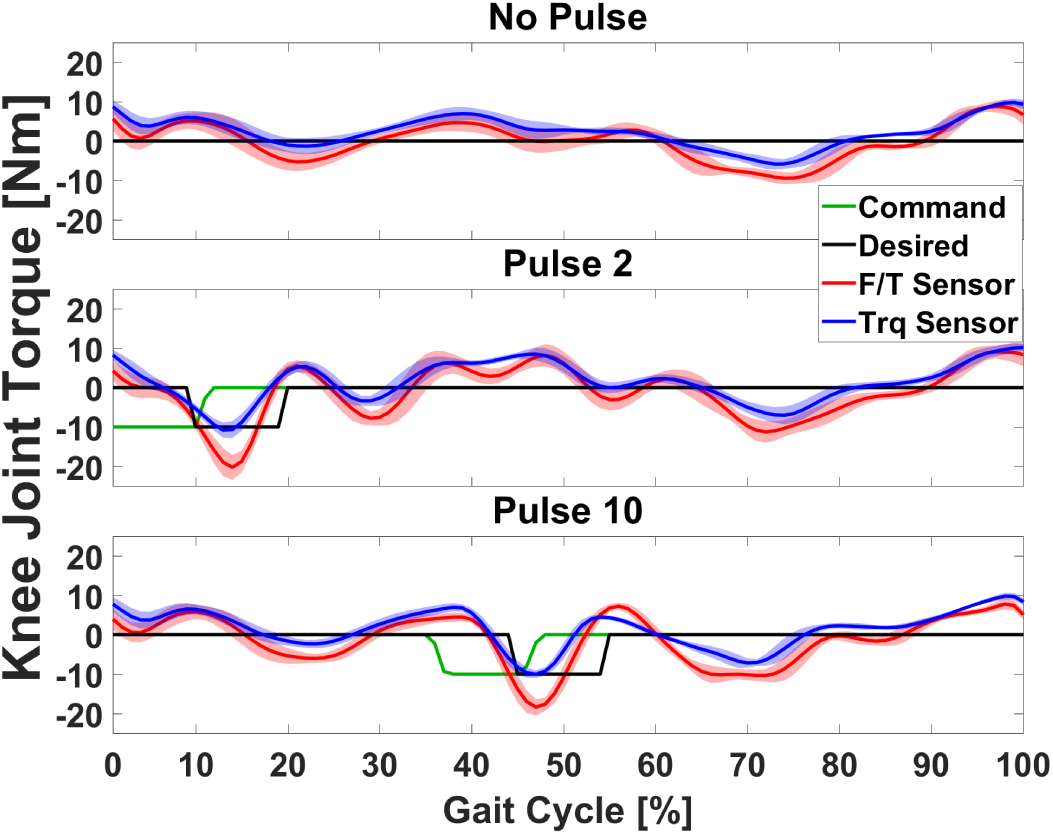
Knee joint torque profiles commanded, desired, estimated based on the 6-axis force/torque sensor and Jacobian transformations, and estimated using the single-axis torque sensor during walking. Estimations account for the expected torque due to gravity such that zero torque corresponds to ideally zero interaction torque. (Top) strides without pulse application, (center) pulse condition two, (bottom) pulse condition ten.

**Fig. S2:**
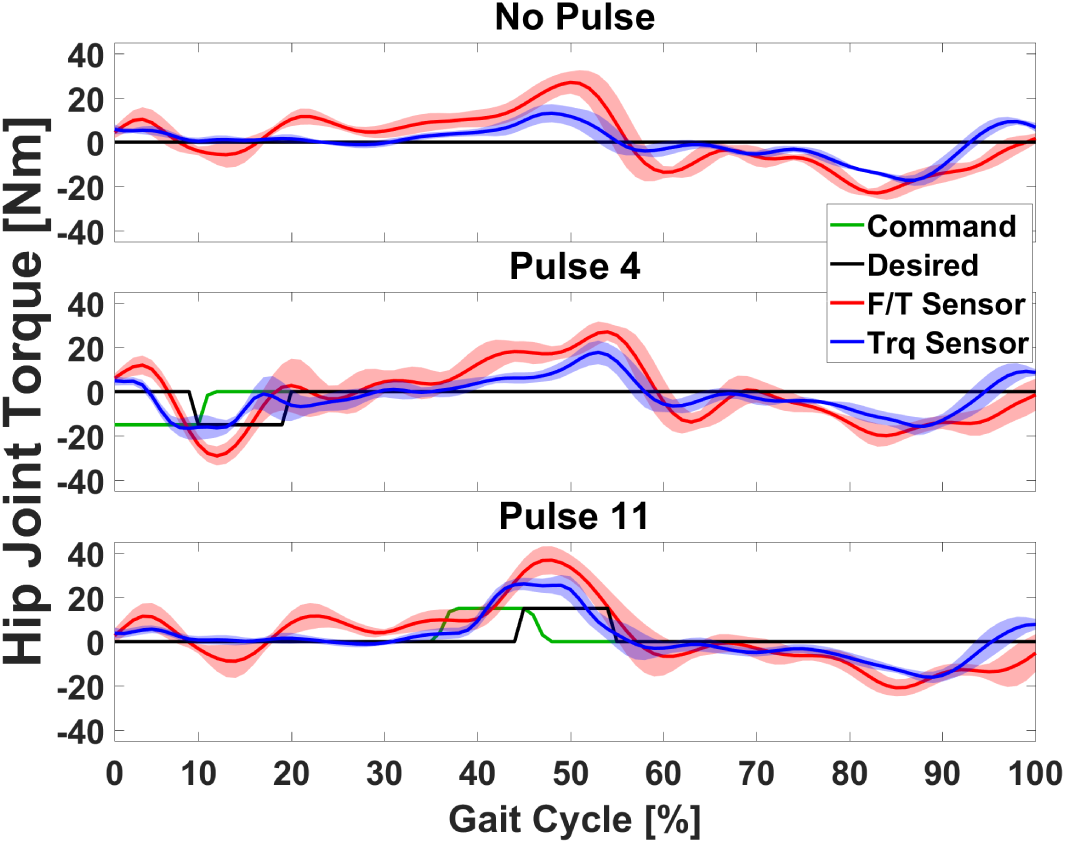
Hip joint torque profiles commanded, desired, estimated based on the 6-axis force/torque sensor and Jacobian transformations, and estimated using the single-axis torque sensor during walking. Estimations account for the expected torque due to gravity such that zero torque corresponds to ideally zero interaction torque. (Top) strides without pulse application, (center) pulse condition two, (bottom) pulse condition ten.

## References

[1] J. Rosen and P. W. Ferguson, eds., Wearable Robotics: Systems and Application. Academic Press, 1st ed., 2020.

[2] B. H. Dobkin and P. W. Duncan, “Should Body WeightSupported Treadmill Training and Robotic-Assistive Steppers for Locomotor Training Trot Back to the Starting Gate?,” Neurorehabil Neural Repair, vol. 26, no. 4, pp. 308–317, 2014.

[3] J. Mehrholz, S. Thomas, C. Werner, J. Kugler, M. Pohl, and B. Elsner, “Electromechanical-assisted training for walking after stroke (Review),” Cochrane Database of Systematic Reviews, no. 5, 2017.

[4] A. Schmid, P. W. Duncan, S. Studenski, S. M. Lai, L. Richards, S. Perera, and S. S. Wu, “Improvements in Speed-Based Gait Classifications Are Meaningful,” Stroke, vol. 38, pp. 2096–2101, 2007.

[5] M. G. Bowden, C. K. Balasubramanian, R. R. Neptune, and S. A. Kautz, “Anterior-Posterior Ground Reaction Forces as a Measure of Paretic Leg Contribution in Hemiparetic Walking,” Stroke, vol. 37, pp. 872–877, 2006.

[6] C. L. Peterson, J. Cheng, S. A. Kautz, and R. R. Neptune, “Leg extension is an important predictor of paretic leg propulsion in hemiparetic walking,” Gait & Posture, vol. 32, no. 4, pp. 451–456, 2010.

[7] H. Hsiao, B. A. Knarr, J. S. Higginson, and S. A. Binder-Macleod, “Mechanisms to increase propulsive force for individuals poststroke.,” Journal of Neuroengineering and Rehabilitation, vol. 12, no. 40, 2015.

[8] H. Hsiao, B. A. Knarr, J. S. Higginson, and S. A. Binder-Macleod, “The Relative Contribution of Ankle Moment and Trailing Limb Angle to Propulsive Force during Gait,” Human Movement Science, pp. 212–221, 2015.

[9] R. L. McGrath, M. Pires-Fernandes, B. Knarr, J. S. Higginson, and F. Sergi, “Toward goal-oriented robotic gait training: The effect of gait speed and stride length on lower extremity joint torques,” IEEE International Conference on Rehabilitation Robotics, pp. 270–275, 2017.

[10] R. L. McGrath, M. L. Ziegler, M. Pires-Fernandes, B. A. Knarr, J. S. Higginson, and F. Sergi, “The effect of stride length on lower extremity joint kinetics at various gait speeds,” PLOS ONE, vol. 14, p. e0200862, feb 2019.

[11] K. N. Winfree, P. Stegall, and S. K. Agrawal, “Design of a minimally constraining, passively supported gait training exoskeleton: ALEX II,” IEEE International Conference on Rehabilitation Robotics, pp. 0–5, 2011.

[12] D. Zanotto, T. Lenzi, P. Stegall, and S. K. Agrawal, “Improving transparency of powered exoskeletons using force/torque sensors on the supporting cuffs,” in IEEE International Conference on Rehabilitation Robotics, pp. 0–5, 2013.

[13] R. L. McGrath and F. Sergi, “Single-stride exposure to pulse torque assistance provided by a robotic exoskeleton at the hip and knee joints,” IEEE International Conference on Rehabilitation Robotics, vol. 2019-June, pp. 874–879, 2019.

[14] N. T. Ray, B. A. Knarr, and J. S. Higginson, “Walking speed changes in response to novel user-driven treadmill control,” Journal of Biomechanics, vol. 78, pp. 143–149, 2018.

